# The Origins of Arginine “Magic”: Guanidinium Like-Charge Ion Pairing and Oligoarginine Aggregation in Water by NMR, Cryoelectron Microscopy, and Molecular Dynamics Simulations

**DOI:** 10.1101/2024.08.04.606526

**Authors:** Denys Biriukov, Zuzana Osifová, Man Nguyen Thi Hong, Philip E. Mason, Martin Dračínský, Pavel Jungwirth, Jan Heyda, Mattia I. Morandi, Mario Vazdar

## Abstract

The phenomenon of like-charge pairing of hydrated ions is a physical manifestation of the unique solvation properties of certain ion pairs in water. Water’s high dielectric constant and related ion screening capability significantly influence the interaction between like-charged ions, with the possibility to transform it – in some cases – from repulsion to attraction. Guanidinium cations (Gdm^+^) represent a quintessential example of such like-charge pairing due to their specific geometry and charge distribution. In this work, we present experimental quantification of Gdm^+^–Gdm^+^ contact ion pairing in water utilizing nuclear magnetic resonance (NMR) spectroscopy experiments complemented by molecular dynamics (MD) simulations and density functional theory (DFT) calculations. The observed interaction is very weak — about –0.5 kJ*·*mol*^−^*^1^ — which aligns with theoretical estimation from MD simulations. We also contrast the behavior of Gdm^+^ with NH_4_^+^ cations, which do no exhibit contact ion pairing in water. DFT calculations predict that the NMR chemical shift of Gdm^+^ dimers is smaller than that of monomers, in agreement with NMR titration curves that display a non-linear Langmuir-like behavior. Additionally, we conducted cryo-electron microscopy experiments on oligoarginines R_9_, which (unlike nona-lysines K_9_) exhibit aggregation in water. This points again to like charge pairing of the guanidinium side chain groups, as corroborated also by molecular dynamics simulations of these peptides in water.

## Introduction

The contact like-charge pairing of ions in water is counterintuitive from the physical point of view since two positively or two negatively charged ions should repel according to Coulomb’s law. However, it has been found that certain like-charge ions do not repel in water but attract each other instead.^1–3^ This qualitative change in ionic interactions is primarily driven by the dielectric constant of the environment (78.5 in water vs. 1 in the gas phase), reducing the electrostatic repulsion forces from *∼*400 kJ*·*mol*^−^*^1^ in the gas phase to only several kJ*·*mol*^−^*^1^ in water. The overall reduction of repulsion is valid for all hydrated ions; however, only in a few cases, repulsion is compensated for by additional attractive van der Waals forces, which qualitatively change the sign of interaction, resulting instead in mutual attraction.^4^ Notably, like-charge ion pairs have been computationally predicted for small spherical ions, *e.g.*, fluoride, by classical molecular dynamics (MD) simulations. The stability of these ion-pairs, however, is attributed to the bridging role of water molecules. Moreover, due to their small size, these ions fit well into the water structure, unlike larger halides.^5^

In contrast, the unique true contact like-charge ion-pairing, without water between the ions, has been computationally predicted for planar, amphiphilic guanidinium (Gdm^+^) cations in numerous MD studies employing both classical force fields and *ab initio* calculations.^3,4,6,7^ Moreover, in the repertoire of like-charged species in water, Gdm^+^–Gdm^+^ pairing is especially important in the biological context since Gdm^+^ is the main group of the essential arginine amino acid side chain. The Gdm^+^ like-charge pairing is observed in many protein structures,^7^ often regulating various protein functions.^8^ However, the experimental evidence of like-charge ion pairs has not been widely reported. Only a few examples currently exist in the literature predicting the Gdm^+^–Gdm^+^ pairing in solution, such as X-ray spectroscopy of Gdm^+^ ions,^9^ mass spectrometry studies of Gdm^+^ water clusters,^10^ as well as electrophoretic mobility experiments of tetraarginines.^11^ Interestingly, the thermodynamic volumetric and activity data of Gdm^+^ salts (which lack the molecular detail) differ significantly from those of salts built of simple spherical ions, indicating anomalies in Gdm^+^ hydration and Gdm^+^– Gdm^+^ interactions.^12,13^ In the biological context, the existence of like-charge aggregates has been suggested by SAXS and NMR experiments for deca-arginines (R_10_) in water.^14^ There is also an indirect indication of nona-arginine (R_9_) aggregation at phospholipid bilayers, as evidenced by fluorescence assays at supported lipid bilayers.^15^

In contrast to rare experimental studies, the estimation of the strength of the weakly interacting like-charge ion pairs is frequently provided by theoretical studies, both at the *ab initio* level^6^ as well as by *ab initio* MD^7^ and classical MD simulations.^3,6,16^ At *ab initio* level, the attraction of the Gdm^+^ ions is observed when dispersion interactions are included, suggesting that the dispersion (van der Waals) forces — unique for Gdm^+^ cation due to its very specific quasiaromatic geometry and electronic structure ^4,6,16^ — play a key role in the total energy balance. Conspicuously, almost all force fields (FFs) used in classical molecular dynamics (MD) simulations predict Gdm^+^–Gdm^+^ like-charge pairing, albeit with different strengths due to favorable van der Waals parameters. In contrast, the interaction between NH^+^ ions (which serve as a mimic of the lysine side chain), as well as lysine-rich peptides, predict the mutual repulsion between them at all levels of theory in all of the above studies. In this work, we focused on two aspects of the like-charge ion pairing by a unique synergistic combination of two experimental techniques supported by theory. First, we applied NMR titration experiments in connection with classical MD simulations and density functional theory (DFT) calculations of NMR shifts, evaluating the experimental free energy of Gdm^+^–Gdm^+^ attraction in water for the first time. Second, we extended the available experimental database of the interaction of arginine-rich peptides in water, providing conclusive evidence of R_9_ aggregation by cryo-electron microscopy (cryo-EM), supported by the classical MD simulations.

## Methods

### Chemicals

Guanidinium chloride, ammonium chloride, sodium chloride, and calcium chloride were purchased from Sigma-Aldrich (St. Louis, MO) with 99 % purity. H-(Lys)_9_-OH (K_9_), and H-(Arg)_9_-OH (R_9_) were synthesized by solid-phase peptide synthesis (SPPS) on PS3 peptide synthesizer (Gyros Protein Technologies, USA) using standard Fmoc chemistry protocols, HBTU coupling reagents, and 2-chlorotrityl chloride resin support (0.1 mmol scale and 10 equivalent amino acid excess). Side-chain-protected peptides were cleaved off the resin with a mixture of acetic acid/2,2,2-trifluoroethanol/dichloromethane (1:1:3) for 2 hours at room temperature. The resin was filtered off, reagents and solvents were evaporated to dryness, and residues were treated with the mixture of water/trifluoroacetic acid (TFA)/triisopropylsilane (95:2.5:2:5). The cleaved and deprotected peptides were lyophilized and purified by RP HPLC (Vydac 218TP101522 column) using methanol and water with 0.05 % TFA as solvents. The purity was assessed by analytical RP-HPLC (Vydac 218TP54 column) and LC/MS (Agilent Technologies 6230 ToF LC/MS). The mass was confirmed by MALDI-ToF MS: H-(Lys)_9_-OH [M+H]^+^ 1171.9; H-(Arg)_9_-OH [M+H]^+^ 1423.9.

### NMR Experiments

In order to study specific ion–ion interactions by NMR, we have eliminated the variation in the mean-field electrostatic screening effects by performing our experiments at a constant ionic strength. We prepared a dilution series with decreasing concentrations of GdmCl and NH_4_Cl, keeping constant ionic strength by adding NaCl in the appropriate amount. The samples were prepared simultaneously to protect them from solvent evaporation, which could negatively affect the salt concentration. 3.1 M and 3.0 M solutions of GdmCl and NH_4_Cl, as well as 6 M GdmCl were prepared in H_2_O/D_2_O mixture (volume ratio 9:1) and diluted by 3 M or 6 M NaCl solution, respectively. For both salt concentrations, chemical shifts were referenced to an internal standard *t* -BuOH (1.24 ppm for ^1^H and 30.29 ppm for ^13^C, respectively). The exception to this is the supplementary measurements in neat water, where chemical shifts were referenced to nithomethane as an external standard (4.40 ppm for ^1^H and 63.23 ppm for ^13^C, respectively). The NMR spectra were recorded on a 500 MHz spectrometer (^1^H at 500 MHz, ^13^C at 125.7 MHz) at room temperature (298 K).

### Cryo-EM Experiments

Cryo-EM samples of concentrated peptide solutions were prepared on either 300 mesh 2/2 Quantifoil or C-flat EM grids (Electron Microscopy Sciences, USA), on which 10 nm protein A-conjugated colloidal gold particles (Au–NP) were preadsorbed (Aurion, Netherlands). Au–NP adsorbed grids were then glow-discharged (30 s, 40 mA) in a Quorum GLOQUBE Plus system. An aliquot (3.0 *µ*L) of either i) 100 mM solution of nona-arginine (R_9_) or nona-lysine (K_9_) in 150 mM NaCl with added 100 mM CaCl_2_ prior to sample freezing or ii) 150 mM NaCl with added 100 mM CaCl_2_ prior to sample freezing was applied to the carbon side of EM grids, and subsequently blotted for 2.0 s at blot force –7 and plunge-frozen into the precooled liquid ethane with a Vitrobot Mark IV (FEI, USA).

Cryo-electron micrographs of vitrified samples were collected using a transmission electron microscope JEOL JEM-2100PLUS (JEOL, Japan) equipped with a TemCam-XF416(ES) (TVIPS, Germany) camera, using an Elsa cryo-transfer holder model 698 (Gatan), at accelerating voltage of 200 keV. Grid mapping and image acquisition were performed using SerialEM software^17^ at a nominal magnification of 80*×* and 2,500*×*, respectively. High-magnification images were recorded at 60,000*×* nominal magnification (0.194 nm pixel size) with a –10*µ*m defocus value. To minimize radiation damage during image acquisition, the low-dose mode in SerialEM software was used, and the electron dose was kept below 100 e*^−^*/Å^2^.

Each high-magnification image of the vitrified samples was first converted to an 8-bit format and binned by 4 to enhance feature recognition. Subsequently, a 250-pixel size square region was selected from each image, the corresponding Fast Fourier Transform (FFT) was calculated using the respective command in Fiji software,^18^ and the normalized radial profile of the FFT power spectra was then measured using Radial Profile Angle plugin of Fiji. In addition, each selected image region was binarized using the center of the gray-level histogram as the threshold, and the resulting spatial autocorrelation function was calculated using a MATLAB script.

### Quantum Chemical Calculations

Density functional theory (DFT) calculations for the carbon-NMR chemical shifts were performed using the single point Gauge-Independent Atomic Orbital (GIAO) method with the B3LYP/aug-cc-pVTZ basis set at the B3LYP-D3/cc-pVDZ optimized geometry (denoted as B3LYP/aug-cc-pVTZ//B3LYP-D3/cc-pVDZ). The calculations were carried out on a Gdm^+^–Gdm^+^ dimer hydrated with 12 water molecules and compared to a Gdm^+^ monomer hydrated with 6 water molecules. Additionally, the levels of theory B3LYP/aug-cc-pVTZ//B3LYP-D3/aug-cc-pVDZ and B3LYP/6-311++G(2d,p)//B3LYP-D3/6-311+G(d,p) were tested for the same system, see Table S1 for a full summary. NH^+^ dimers — prepared as hydrated monomers with 4 water molecules per each and calculated at the same level of theory — were found unstable and dissociated, resulting in a formation of solvent-separated ion-pair instead, Figure S1. Calculated carbon-NMR shifts were subtracted from the reference carbon-NMR shift tetra-methyl silane (TMS) at the same level of theory. Additionally, carbon-NMR shift calculations were performed with the polarizable continuum model (PCM) at B3LYP/aug-cc-pVTZ//B3LYP-D3/aug-cc-pVDZ level of theory for Gdm^+^–Gdm^+^ dimers.^19,20^ The carbon-NMR shifts were also calculated for the Gdm^+^–Gdm^+^ dimers with the central carbon distance from 2.9 Å to 5.5 Å in order to determine the cut-off distance beyond which the carbon-NMR shift remains relatively constant, Figure S2. The Grimme empirical dispersion correction was used for geometry optimizations of microhydrated clusters at the B3LYP level of theory.^21^ All DFT calculations are performed with the Gaussian16 suite of codes.^22^

### Molecular Dynamics Simulations

All-atom MD simulations were designed to resemble the composition of aqueous solutions used in the NMR and cryo-EM experiments and provide direct atomistic insights into the studied phenomena.

To resemble NMR systems, 400 GdmCl and 330 NH_4_Cl ion pairs were dissolved in 5550 water molecules to achieve the desired concentration of *∼*3.0 M. Systems with lower GdmCl and NH_4_Cl concentrations were prepared by adjusting the amount of Gdm^+^ to [10, 25, 50, 100, 200] or NH^+^ to [10, 20, 40, 80, 160] and adding the corresponding amount of sodium cations to have a total of 400 and 330 ion pairs, respectively. Similarly, 1100 GdmCl ion pairs were dissolved in 5500 water molecules to achieve *∼*6.0 M concentration, and [50, 100, 200, 400, 800] Gdm^+^ with added sodium were used to create lower concentrations. Thus, the total ionic strength of the solutions was always kept constant as in the experiments. Three different FFs were tested for GdmCl: Kirkwood-Buff (KB) FF,^23,24^ which does not predict Gdm^+^–Gdm^+^ pairing; the updated version of KB model (KB2) that introduces Gdm^+^– Gdm^+^ pairing into the original model;^25^ and prosECCo75 model^26^ — the CHARMM36^27^ derivative that incorporates the electronic polarization via charge scaling ^28^ and also suggests Gdm^+^–Gdm^+^ pairing. For NH_4_Cl simulations, only prosECCo75 model was used.

To compare the simulations with cryo-EM analysis, an additional set of MD simulations was conducted for aqueous solutions of R_9_ and K_9_ peptides. Two peptide concentrations were examined under various ionic strengths. Specifically, 6 and 16 peptide molecules (R_9_ or K_9_), corresponding to concentrations of 0.02 M and 0.05 M, were solvated in aqueous solutions containing either no salt (apart from chloride counterions necessary to neutralize the positive net charge of the peptides), 0.13 M NaCl, or 0.13 M CaCl_2_. All simulation systems contained 15856 water molecules. The prosECCo75 FF was employed for these simulations.

All MD simulations were performed using a leap-frog integrator with a time step of 2 fs in GROMACS engine.^29^ We adopted simulation protocols in accord with previously used and recommended for the corresponding FFs. Buffered Verlet lists^30^ were used to keep track of atomic neighbors. The cutoff for short-range electrostatic interactions was set to 1.2 nm (prosECCo75) or 1 nm (KB and KB2). Long-range electrostatics was treated using smooth particle mesh Ewald (PME) algorithm.^31^ The cutoff for Lennard–Jones interactions was set to 1.2 nm with the forces switched to zero starting at a distance of 1.0 nm ^32^ (prosECCo75) or simply set to 1.0 nm (KB and KB2) The temperatures of 300 K (for the GdmCl and NH_4_Cl solutions) and 310 K (for R_9_ and K_9_ solutions) were maintained by the Nośe–Hoover thermostat^33,34^ with a coupling time constant of 1 ps (prosECCo75) or v-rescale thermostat^35^ with a coupling time of 0.1 ps (KB and KB2). The Parrinello–Rahman barostat^36^ with a coupling time constant of 5 ps (prosECCo75) or 2 ps (KB and KB2) and compressibility of 4.5*×*10*^−^*^5^ bar*^−^*^1^ was maintaining the pressure of 1 bar. Water was simulated as a CHARMM-specific TIP3P model^37,38^ (prosECCo75) or SPC/E model^39^ (KB and KB2). Bonds in solute molecules involving hydrogen atoms were constrained using P-LINCS, ^40,41^ whereas the geometry of water was constrained using SETTLE.^42^ In the case of KB simulations, all other covalent bonds were also constrained, following the original simulation protocol.^24^ Simulations were 500 ns long, and the first 100 ns were omitted from all analyses.

## Results and Discussion

### Like-Charge Pairing of Gdm**^+^** Ions in Water

Figure 1A shows the experimental NMR chemical shifts with increasing Gdm^+^ or NH^+^ concentrations in GdmCl and NH_4_Cl solutions, respectively. The experiments were performed at a constant ionic strength (*i.e.*, at constant mean-field electrostatic screening of studied solutions) of approximately 3 M, which was necessary for accurate free energy determination of ion-specific Gdm^+^–Gdm^+^ pairing. When experiments are conducted in neat water, the increase of GdmCl concentration changes the ^1^H NMR and ^13^C NMR shifts of the Gdm^+^ proton and carbon atoms, respectively, as illustrated in Figure S3 in the Supporting Information (SI). The approach of Gdm^+^/Na^+^ titration has previously been demonstrated to be successful in measuring the electrophoretic mobility of GdmCl solutions.^11^

The nonlinear change in the observed ^1^H NMR chemical shift indicates cation–cation binding, as shown for GdmCl solutions. The measured difference Δ*δ* in ^1^H NMR shift can be used to characterize and quantify the energetics of binding by calculating the dissociation constant *K_d_*.^43^ The dependence of Δ*δ* on Gdm^+^ concentration follows a typical Langmuir isotherm, which can be fitted as:

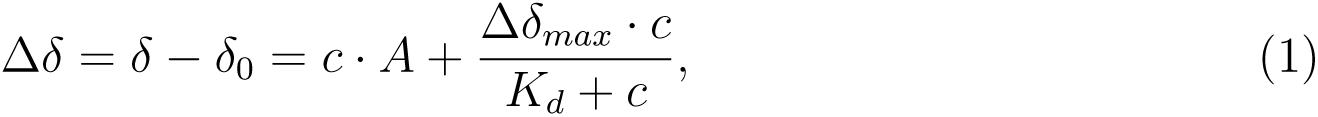

**Figure 1:**
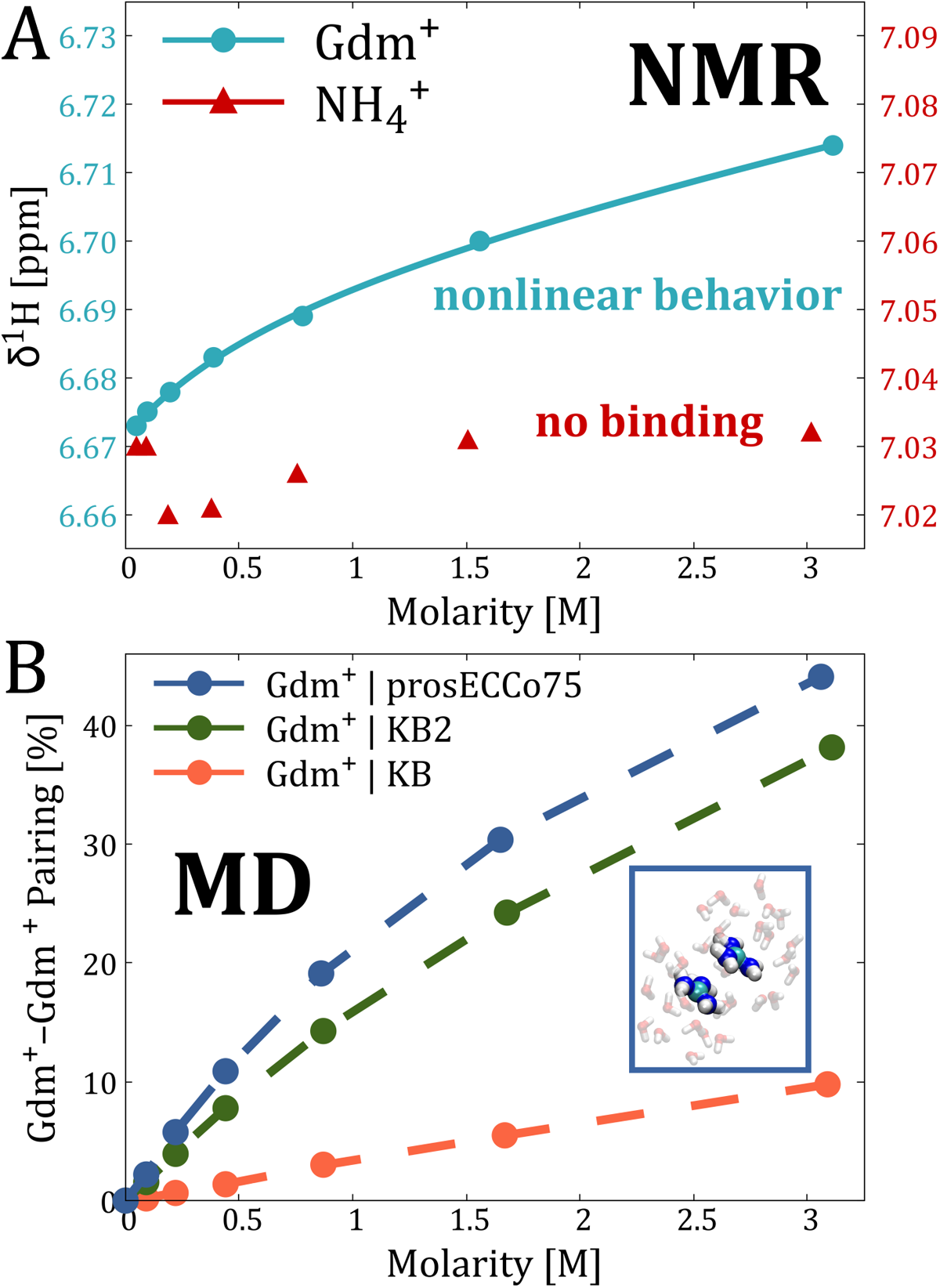
**A)** The concentration dependence of ^1^H NMR chemical shift on the Gdm^+^ or NH_4_^+^ concentration in an aqueous solution, where a constant ionic strength of *∼*3 M is maintained by adding NaCl. **B)** The like-charge pairing of the Gdm^+^ cations as a function of Gdm^+^ concentration in an aqueous solution, where a constant ionic strength of *∼*3 M is maintained by adding NaCl. Data for the three MD force fields are compared: prosECCo75 and KB2 models show nonlinear behavior and, at the same time, capture contact ion Gdm^+^–Gdm^+^ pairing, while KB model shows the linear behavior and does not capture contact ion Gdm^+^–Gdm^+^ pairing. The blue inset rectangle exemplifies Gdm^+^–Gdm^+^ contact ion pairing from MD simulations using the prosECCo75 force field.

where *A* is a linear constant with units of chemical shift per molar concentration and Δ*δ_max_* is the maximum change in chemical shift observed for the nonlinear portion of the curve at saturation.

We calculated the dissociation constant *K_d_* for 3 M GdmCl solution to be *∼*0.8. This value can be converted to the free energy of binding using the formula Δ*G_b_* = *−kT* ln *K_b_*, where *K_b_* is the binding constant, which is the inverse of the dissociation constant *K_d_*. Using this approach, we calculated the binding free energy of Gdm^+^ ions to be approximately –0.2 kT or –0.5 kJ*·*mol*^−^*^1^ at room temperature. Thus, Gdm^+^ ions form a weakly stabilized contact ion pair in a solution of 3 M ionic strength. This finding provides the first experimental evaluation of Gdm^+^–Gdm^+^ contact ion pairing strength in aqueous solutions. In the solution of 6 M ionic strength, the like-charge pairing is stronger, which is expected since the higher ionic strength further reduces electrostatic repulsion. This behavior is illustrated in Figure S4, where a non-linear increase in ^1^H NMR shift leads to a Δ*G_b_* of –1.2 kT or –3.0 kJ*·*mol*^−^*^1^.

Finally, as a negative control, we examined the NH_4_Cl system where MD simulations predict no like-charge pairing.^6,7^ The measured ^1^H NMR chemical shifts for the NH_4_Cl solution at a constant ionic strength of 3 M exhibit no discernible trend, Figure 1A. This behavior is qualitatively different from the GdmCl experiments, indicating the absence of attraction for NH^+^ ions in water.

Since the experimental NMR measurements cannot provide all the molecular details of the like-charge ion pairing, we performed MD simulations of the investigated systems. We modeled systems with varying concentrations of GdmCl and NH_4_Cl, maintaining constant ionic strengths of approximately 3 M and 6 M by adding the appropriate amount of NaCl (see Methods for details). For the simulations involving Gdm^+^, we utilized three different FFs: KB, KB2, and prosECCo75. For systems containing NH^+^, we used the prosECCo75. Ion pairing was quantified using two different distance cutoffs. The first cutoff was set at 3.9 Å for C*_z_*–C*_z_* and 4.9 Å for N*_z_*–N*_z_* in GdmCl and NH_4_Cl, respectively. These values represent the maximum in the corresponding radial distribution functions calculated from the MD simulations, Figure S5. The second cutoff of 4.5 Å was applied to both systems and derived from NMR chemical shift PCM/GIAO calculations on the Gdm^+^–Gdm^+^ dimer (see Methods). The calculated ^13^C NMR shift does not significantly change beyond a 4.5 Å distance between guanidinium carbons, see Figure S2. Therefore, we presume that beyond this cutoff, the Gdm^+^ cations do not “feel” each other in the NMR experiments. This observation can also be extended to the interpretation of ^1^H NMR experiments.^44^

Our MD data, summarized in Figure 1B, show the calculated contact probabilities as a function of Gdm^+^ concentration in GdmCl solutions at a constant ionic strength of *∼*3 M. Notably, two of the three FFs examined — KB2 and prosECCo75 — exhibit a nonlinear trend that aligns with the NMR experiments. This trend indicates significant binding between Gdm^+^ cations, with prosECCo75 showing slightly stronger interactions. Importantly, only these two FFs capture the contact ion pairing of Gdm^+^ cations, as shown in the inset of Figure 1B and the representative radial distribution functions in Figure S5. In contrast, the KB model demonstrates a linear trend, which indicates overall weak, solvent-shared binding between Gdm^+^ cations. Similarly, the data for NH_4_Cl solutions reveal an exclusively linear trend with increasing NH^+^ concentration, Figure S7. All these conclusions hold true across both concentrations and cutoffs used to distinguish contact ion pairing, Figures S7 and S8. The radial distribution functions shown in Figure S5 can also be converted into the potential of mean force, which provides a rough estimate of the binding free energy Δ*G_b_*between a pair of Gdm^+^ ions, Figure S6. Using RDFs derived from MD simulations of *∼*3 M GdmCl solution, the estimated Δ*G_b_* from the energy minima is found to be –1.1 and –0.6 kJ*·*mol*^−^*^1^ for the prosECCo75 and KB2 models, respectively, Table S2, which is in excellent agreement with the experimental data.

We also compared the experimental NMR results with DFT calculations of GIAO NMR chemical shifts for optimized microhydrated Gdm^+^ clusters, Figure S1. Specifically, we calculated the ^13^C NMR chemical shift for a Gdm^+^–Gdm^+^ dimer hydrated with 12 water molecules and compared it to a Gdm^+^ monomer hydrated with 6 water molecules at various levels of theory (see Methods). In all cases, the ^13^C NMR shift of the dimer was consistently lower than that of the monomer, with differences ranging from 0.8 to 1.4 ppm, Table S1. Additionally, PCM calculations for Gdm^+^ ions without hydrating water showed a reduced shift difference of 0.3 ppm. The observation that the ^13^C NMR chemical shift is lower in the dimer compared to the monomer is also experimentally validated. ^13^C NMR experiments for GdmCl in 6 M ionic strength solutions demonstrate a nonlinear behavior of the chemical shift with increasing GdmCl concentration, similar to the ^1^H NMR experiments, as shown in Figure S4. The estimated Δ*G_b_* from the ^13^C NMR data is –3.7 kJ*·*mol*^−^*^1^, which agrees well with the ^1^H NMR data.

### Like-Charge Pairing of Oligoarginines in Water

To further experimentally assess guanidinium-based ion pairing, we evaluated the aggregation of biologically relevant oligopeptides in water using cryo-EM measurements. We compared the behavior of Gdm^+^-containing peptides (nona-arginine, R_9_) with NH^+^-based ones (nona-lysine, K_9_). Specifically, R_9_ peptides are widely used as cell-penetrating peptides, and their aggregation properties in bulk might be relevant for their translocation efficiency.^15,45^

Conventionally, cryo-EM is utilized to resolve protein structures or to image self-assembling systems such as lipid membranes and organic or metallic nanoparticles. However, it can also visualize solutions of nanoscopic molecules, ^46^ as demonstrated with heavy atom nanocrystals^47^ and surfactant micelles.^48^ Images of unstructured systems, such as peptides in solution, often exhibit low contrast due to the small size and composition differences compared to the surrounding bulk amorphous ice, ^48,49^ which limits the obtainable information. Here, to enhance the contrast of our systems and thus effectively assess differences in peptide aggregation, we modulated the microscope Contrast Transfer Function (CTF) by shifting the image defocus to large values (–10 *µ*m). By acquiring images at high defocus values, the signal from low spatial frequencies (lower resolution information) is increased, thus enhancing the contrast.^50,51^ This approach allows us to visualize features that would otherwise be difficult to identify. However, it should be noted that high-frequency information is significantly reduced with this method.

To maintain proper vitrification of the samples into glassy ice, the peptide concentration was maintained at 100 mM, but aggregation and clustering were enhanced by increasing the ionic strength of the buffer via the addition of 100 mM CaCl_2_. The cryo-EM images of the two peptide solutions display distinct patterns and features, visible in both grayscale and binarized images, see Figure 2. The R_9_ peptide solution displays characteristic spatial clustering of the signal, while the K_9_ peptide solution possesses negligible features or clusters. The spatial clustering of similar gray-value pixels is consistent with previously reported images of surfactant micelles,^48^ suggesting the presence of nanometric (or potentially sub-nanometric) objects in the solution. The two-dimensional FFT power spectra of the respective images further underscore the differences between the two peptides. The R_9_ peptide solution displays a strong low-frequency signal and a first Thon ring,^52^ characteristic of significant contrasted features within the images. Conversely, the K_9_ peptide solution shows a lower intensity in the low spatial frequency region, indicative of fewer features. Comparing these images with those obtained from buffer-only samples (150 mM NaCl and 100 mM CaCl_9_) reveals that the signal from the K_9_ peptide solution is closer to that of a peptide-free solution, Figure S9.

The analysis of the radial profile of the respective FFT power spectra confirms that R_9_ displays an enhanced signal in the 0.09–0.2 nm*^−^*^1^ spatial frequency region, with an additional peak located at 0.4 nm*^−^*^1^ (Figure 3A). In contrast, the FFT of K_9_ images show only a minor increase in signal at 0.15 nm*^−^*^1^ without any additional Thon rings present. When compared to the FFT of buffer-only images, it is evident that the K_9_ power spectra mostly overlap with peptide-free samples, Figure S9C, suggesting the absence of significant features in these images. Conversely, the distinct first and second peaks in the R_9_, which are absent in buffer-only solutions (Figure S9B), indicate that the observed spatial clustering is a unique characteristic of the R_9_ images.

**Figure 2:**
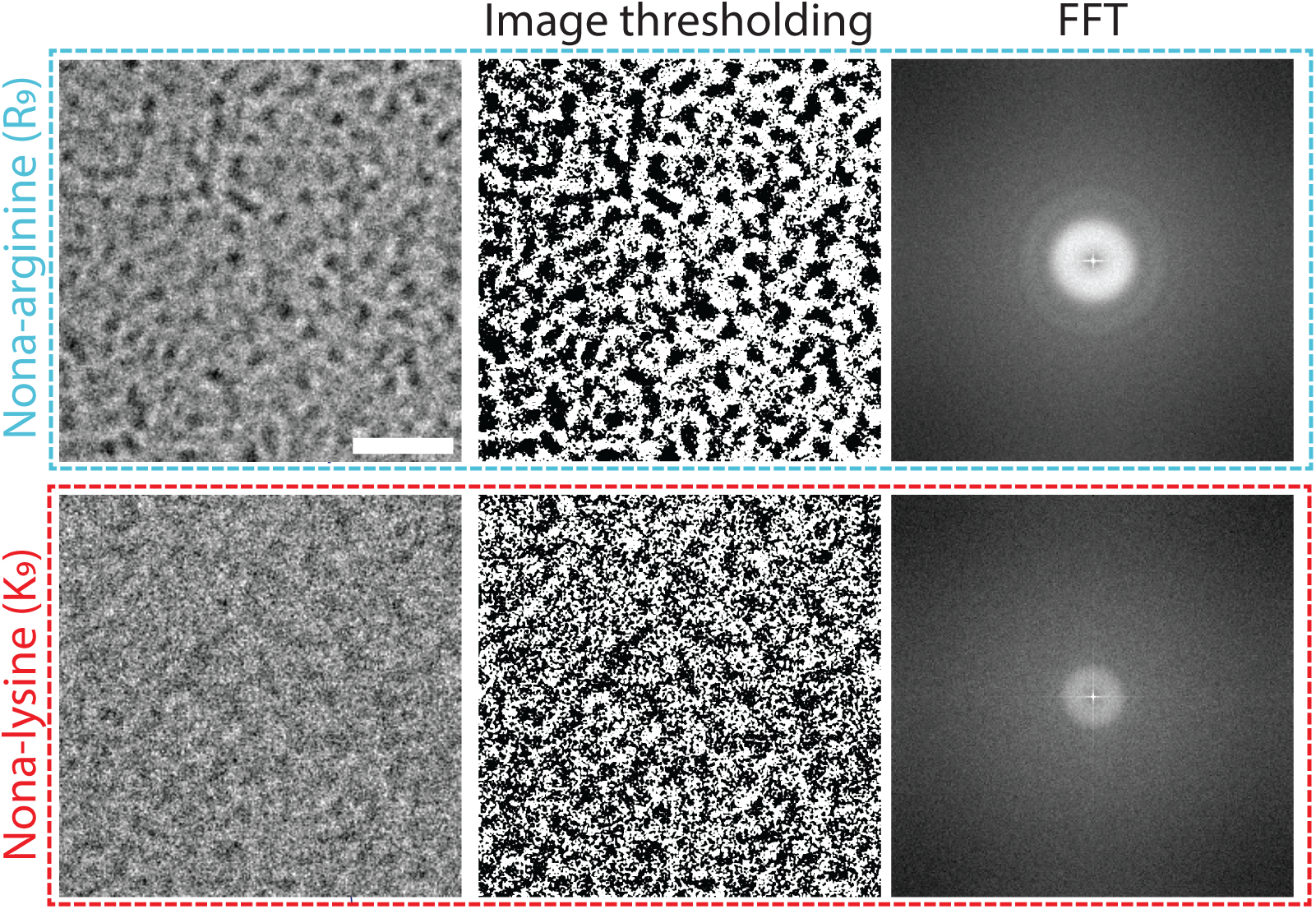
Representative cryo-EM images of 100 mM R_9_ (top, blue) and K_9_ (bottom, red) in 150 mM NaCl and 100 mM CaCl_2_ buffer, along with their corresponding binarized images used for autocorrelation function calculations. The images display distinct differences in the spatial distribution of the intensity signals. The R_9_ peptide solution exhibits clusters of pixels with the same gray values, while the K_9_ peptide solution has a more diffuse distribution. Calculation of the median FFT from all replicates (n = 205 for R_9_ and n = 152 for K_9_) confirms significant differences between the two peptides. The scale bar for the cryo-EM images is 50 nm.

Additionally, analyzing the spatial autocorrelation function (ACF) of the images further supports these findings, Figure 3B. The gray-value correlations for K_9_ rapidly decay to no correlation after three neighboring pixels, with minimal anticorrelation (*i.e.*, negative ACF values). In contrast, R_9_ signal remains correlated for approximately five pixels and subsequently exhibits strong anticorrelation (opposite gray values). Comparing the ACF curves of the peptides solutions with those of the buffer-only images further confirms that K_9_ more closely resembles a peptide-free scenario, Figure S9C and S9D. While our analysis does not enable precise determination of cluster sizes due to resolution loss caused by high defocus during acquisition, it nevertheless demonstrates that the presence of Gdm^+^ moieties in the peptides significantly promotes aggregation/clustering compared to NH^+^ moieties.

**Figure 3:**
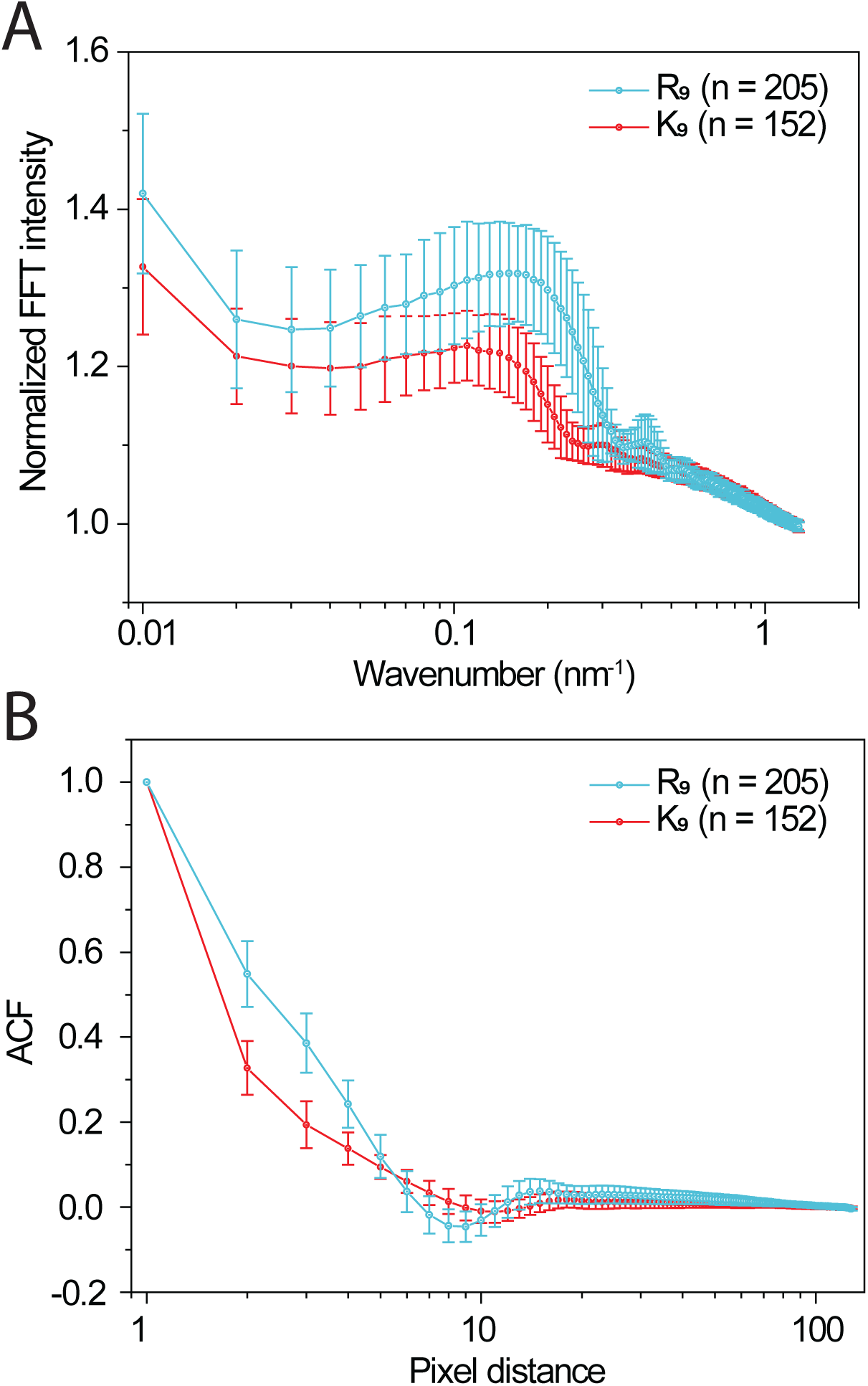
(A) Radial profiles derived from FFT power spectra images that show a significantly different spatial signal distribution for R_9_ (blue) compared to K_9_ (red). (B) Spatial autocorrelation function curves calculated from cryo-EM images, representing the correlation of grayscale values as a function of pixel distance from each pixel. For R_9_ (blue), the data reveal strong positive correlation (same gray values) at short pixel distances, followed by pronounced anticorrelation (opposite gray values), and ultimately a stochastic distribution at longer distances (tail baseline), altogether suggesting the presence of clusters. In contrast, the K_9_ (red) data show a rapid decay to stochastic values, indicating minimal spatial correlation. Data are presented as mean *±* standard deviation, based on n = 205 frames for R_9_ and n = 152 frames for K_9_.

To further validate the cryo-EM findings, we conducted molecular dynamics simulations of R_9_ and K_9_ peptides at two different concentrations (around 0.02 M and 0.05 M) in aqueous solutions containing either counterions only or NaCl/CaCl_2_ at a concentration of *∼*0.13 M. The aggregation behavior of R_9_ and K_9_ peptides was analyzed using radial distribution functions (RDFs, *g*(*r*)) for the C*_z_*–C*_z_* (Gdm^+^) and N*_z_*–N*_z_* (NH^+^) atom pairs, respectively. As shown in Figure 4, the RDFs excluding intramolecular contacts of C*_z_*–C*_z_* and N*_z_*–N*_z_* display distinct patterns. The *g*_Cz_*_−_*_Cz_ (*r*) functions exhibit sharp first peaks at a distance of approximately 0.39 nm across all studied systems, indicating significant interactions between Gdm^+^ groups of arginine residues in different R_9_ molecules, *i.e.*, not within the same peptide. In contrast, the *g*_Nz_*_−_*_Nz_(*r*) functions do not show any clear peaks, suggesting a lack of similar interactions between NH^+^ groups in K_9_ peptides. Notably, the interactions between Gdm^+^ groups is enhanced by increasing ionic strength or peptide concentration. However, in the presence of CaCl_2_ and at lower peptide concentration, the intensity of the first peak is higher compared to higher peptide concentration, indicating stronger aggregation at lower peptide concentrations.

## Conclusions

In this work, we performed NMR titration experiments on aqueous solutions of GdmCl and NH_4_Cl, maintaining constant ionic strength by adding appropriate amounts of NaCl. For the first time, we experimentally quantified the strength of Gdm^+^–Gdm^+^ like-charge ion pairing, which is not observed with NH_4_^+^ ions. In particular, the pairing strength is approximately –0.5 kJ*·*mol*^−^*^1^ in a solution with an ionic strength of 3 M. Our experimental findings are in excellent agreement with theoretical predictions from MD simulations and are further corroborated by results from DFT calculations. Next, we performed cryo-EM experiments on nona-arginines (R_9_) and nona-lysines (K_9_) peptides in aqueous solutions containing NaCl and CaCl_2_. We conclusively demonstrated that R_9_ peptides aggregate in water, unlike K_9_ peptides. This finding was further confirmed by MD simulations of solvated nona-peptides, where only R_9_ peptides showed a tendency to aggregate. Overall, our combined use of NMR and cryo-EM techniques, alongside theoretical methods, reveals that Gdm^+^-containing species exhibit counterintuitive like-charge ion pairing. This finding underscores the distinctive physicochemical behavior of Gdm^+^ ions, which is driven by their specific chemical properties, unique electronic structure, and water solvation.

**Figure 4:**
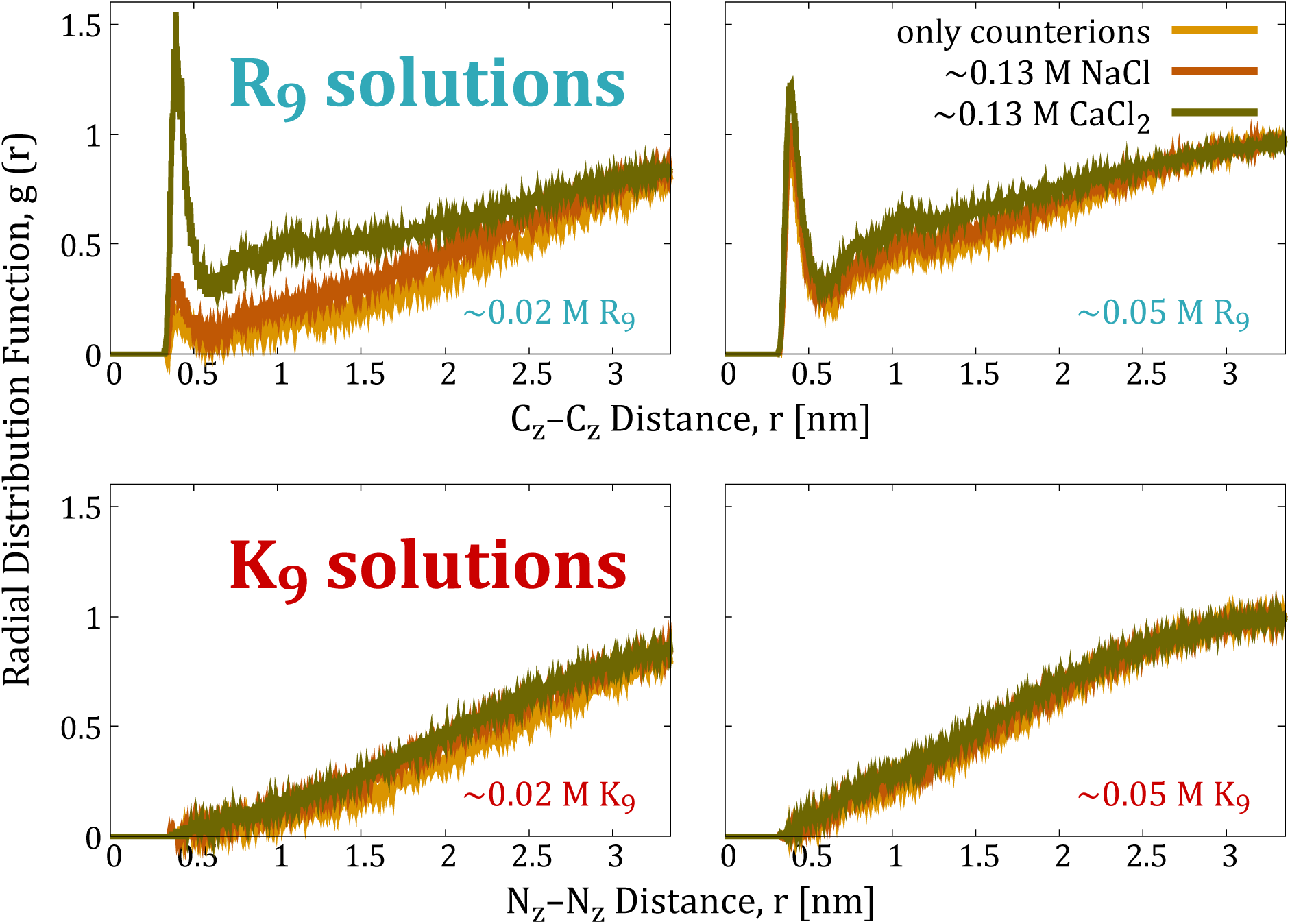
Radial distribution functions for C*_z_*–C*_z_* (in Gdm^+^ groups) and N*_z_*–N*_z_* (in NH_4_^+^ groups) calculated from MD simulations of R_9_ or K_9_ in various aqueous solutions. Note that intramolecular interactions within a single peptide are excluded from these RDFs.

## Acknowledgement

D.B. acknowledges support from the project “National Institute of Virology and Bacteriology (Program EXCELES, ID Project No. LX22NPO5103) – Funded by the European Union – Next Generation EU”. P.J. acknowledges support from the European Research Council via an ERC Advanced Grant no. 101095957. M.V. and J.H. acknowledge support by the project “The Energy Conversion and Storage”, funded as project No. CZ.02.01.01/00/22 008/0004617 by Programme Johannes Amos Commenius, call Excellent Research. M.V. acknowledges support by the Ministry of Education, Youth and Sports of the Czech Republic through the e-INFRA CZ (ID:90254), Project OPEN-30-53. The authors would like to acknowledge the contribution of COST Action CA21169, supported by COST (European Cooperation in Science and Technology). The authors would like to thank the group of Tomáš Kouba for the assistance with cryo-EM acquisition and sample preparation. The authors also acknowledge Grammarly and ChatGPT for improving the readability and language of the manuscript.

---

**TOC Graphic.**
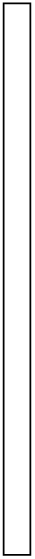

## Notes

### Competing Interest Statement

The authors have declared no competing interest.

