## Supporting Information for "The Origins of Arginine “Magic”: Guanidinium Like-Charge Ion Pairing and Oligoarginine Aggregation in Water by NMR, Cryoelectron Microscopy, and Molecular Dynamics Simulations"

### Quantum Mechanical Calculations

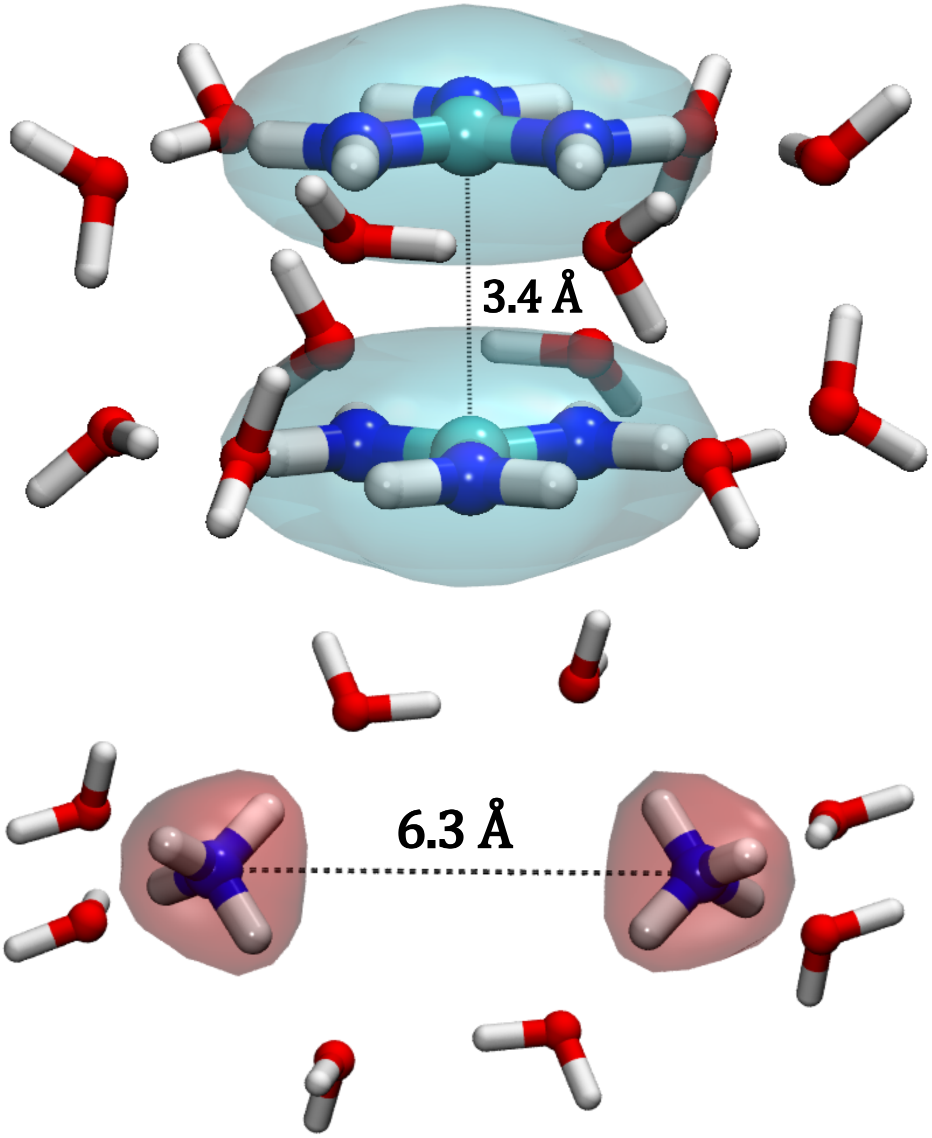


**Figure S1:** Optimized $Gdm^{+}$–$Gdm^{+}$ (top) and $NH_4^{+}$–$NH_4^{+}$ (bottom) dimers at B3LYP-D3-aug-cc-pVDZ level of theory. $Gdm^{+}$ and $NH_4^{+}$ ions are depicted in blue and red colors, respectively, using QuickSurf visualization in VMD,[S1] with these colors corresponding to the data lines in other figures for consistency.

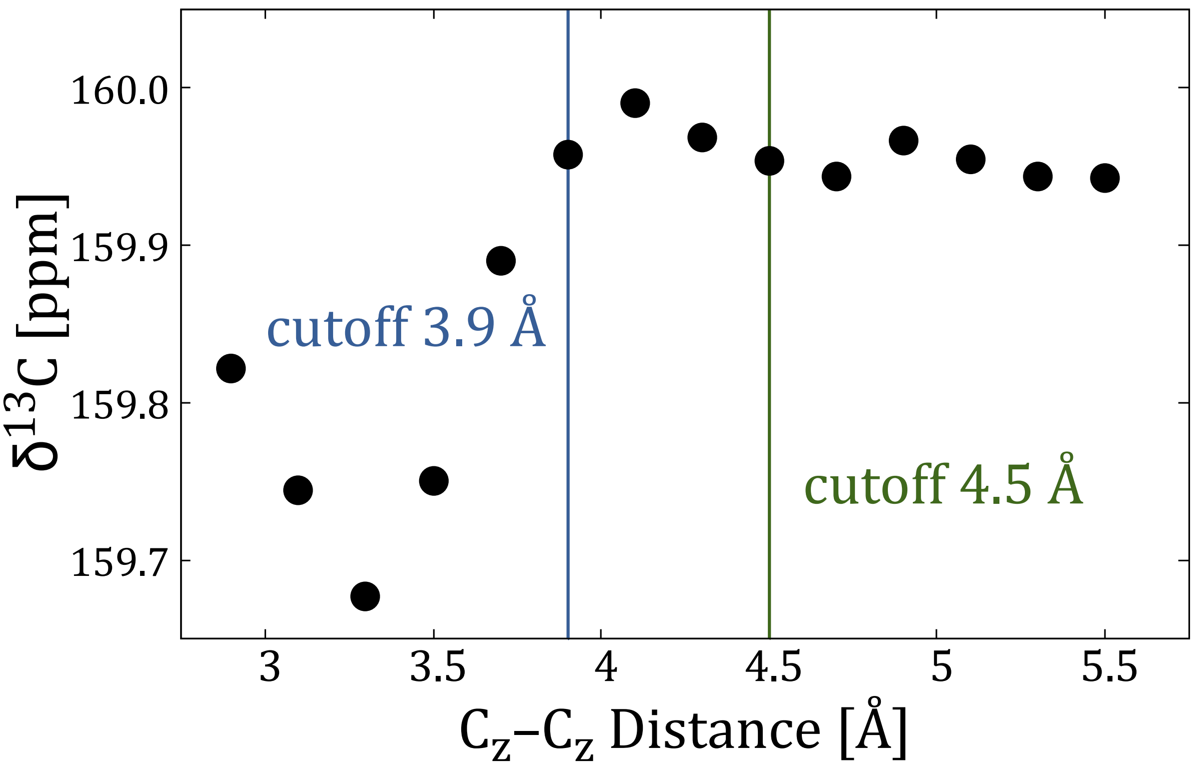


**Figure S2:** The dependence of the $^{13}$C-NMR chemical shift for the Gdm$^+$–Gdm$^+$ dimer on the distance between guanidinium carbons. The data are calculated from quantum chemical calculations at the PCM/B3LYP/aug-cc-pVTZ//B3LYP-D3-aug-cc-pVDZ level of accuracy.

**Table S1:** Predicted $^{13}$C-NMR chemical shifts for the Gdm$^+$–Gdm$^+$ dimer and Gdm$^+$ monomer, along with the corresponding differences, calculated using quantum mechanical methods with different levels of accuracy and solvent representations.

| Method | Monomer [ppm] | Dimer [ppm] | Δ [ppm] |
|---:|---:|---:|---:|
| B3LYP/aug-cc-pVTZ//B3LYP-D3-cc-pVDZ | 160.19 | 158.78 | 1.41 |
| B3LYP/aug-cc-pVTZ//B3LYP-D3-aug-cc-pVDZ | 160.17 | 159.40 | 0.77 |
| B3LYP/6-311++G(2d,p)//B3LYP-D3-6-31+G(d) | 160.47 | 159.62 | 0.85 |
| PCM/B3LYP/aug-cc-pVTZ//B3LYP-D3-aug-cc-pVDZ | 160.01 | 159.69 | 0.32 |

### NMR Experiments

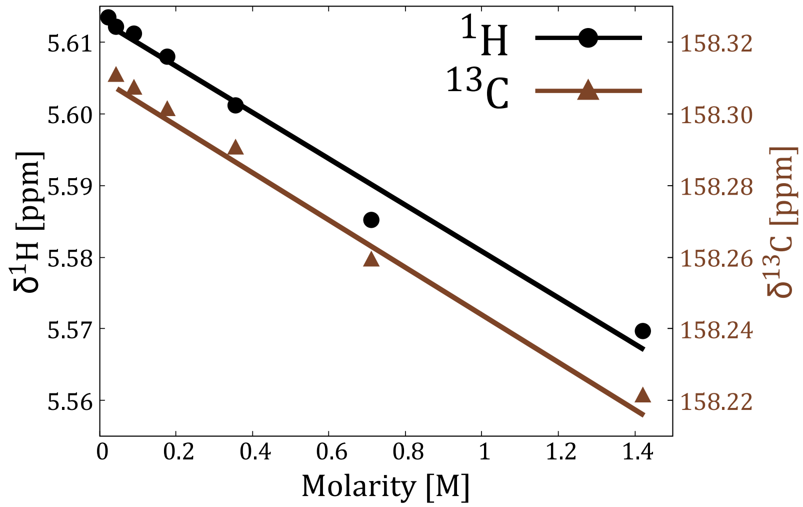


**Figure S3:** $^1$H-NMR and $^{13}$C-NMR shifts of GdmCl solution in neat water.

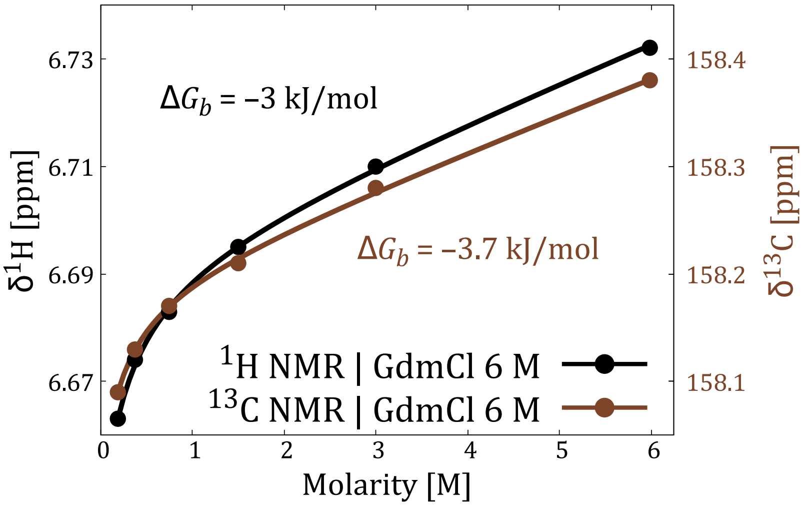


**Figure S4:** $^1$H-NMR and $^{13}$C-NMR shifts of GdmCl solution in the solution with 6 M ionic strength.

### Molecular Dynamics Simulations

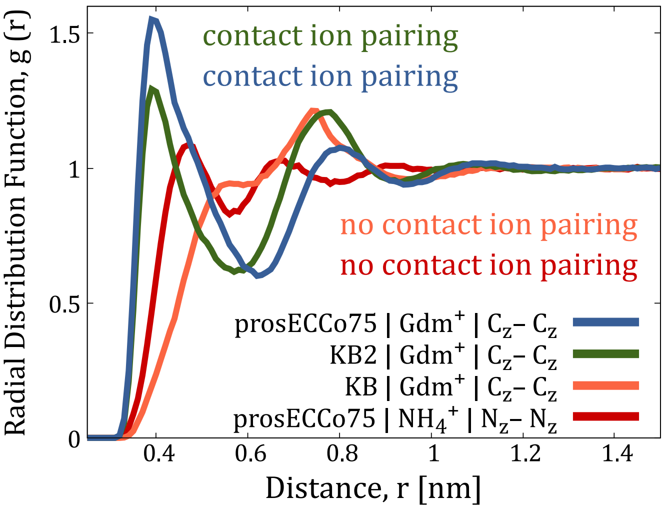


**Figure S5:** Radial distribution functions from MD simulations of GdmCl and $NH_4Cl$ solutions at a constant ionic strength of ∼3 M.

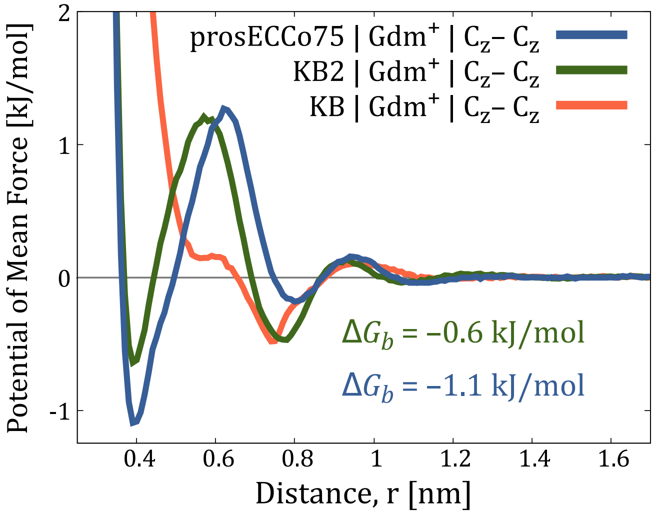


**Figure S6:** Potential of mean force calculated from the radial distribution functions from MD simulations of GdmCl solutions at a constant ionic strength of ∼3 M.

**Table S2:** Estimation of $Gdm^+$–$Gdm^+$ binding free energy $\Delta G_b$ from MD simulations using different force fields and varying the total ionic strength and $Gdm^+$ concentration.

| Force Field | Total Concentration [M] | $Gdm^+$ Concentration [M] | $\Delta G_b$ [kJ·mol$^{-1}$] |
|---|---|---|---|
| prosECCo75 | ∼3M | ∼0.1M | –2.5 |
| | | ∼3M | –1.1 |
| | ∼6M | ∼0.5M | –3.1 |
| | | ∼6M | –0.6 |
| KB2 | ∼3M | ∼0.1M | –1.4 |
| | | ∼3M | –0.6 |
| | ∼6M | ∼0.5M | –1.7 |
| | | ∼6M | –0.4 |

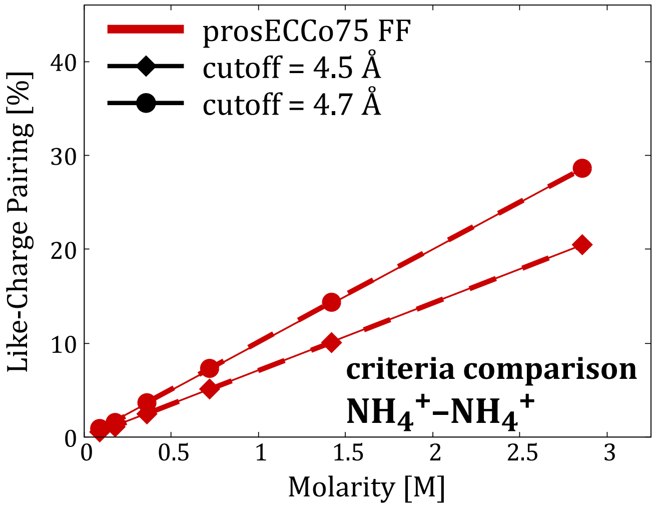


**Figure S7:** The ion pairing of the $NH_4^+$ cations as a function of $NH_4^+$ concentration in an aqueous solution, where a constant ionic concentration of ∼3 M is maintained by adding NaCl.

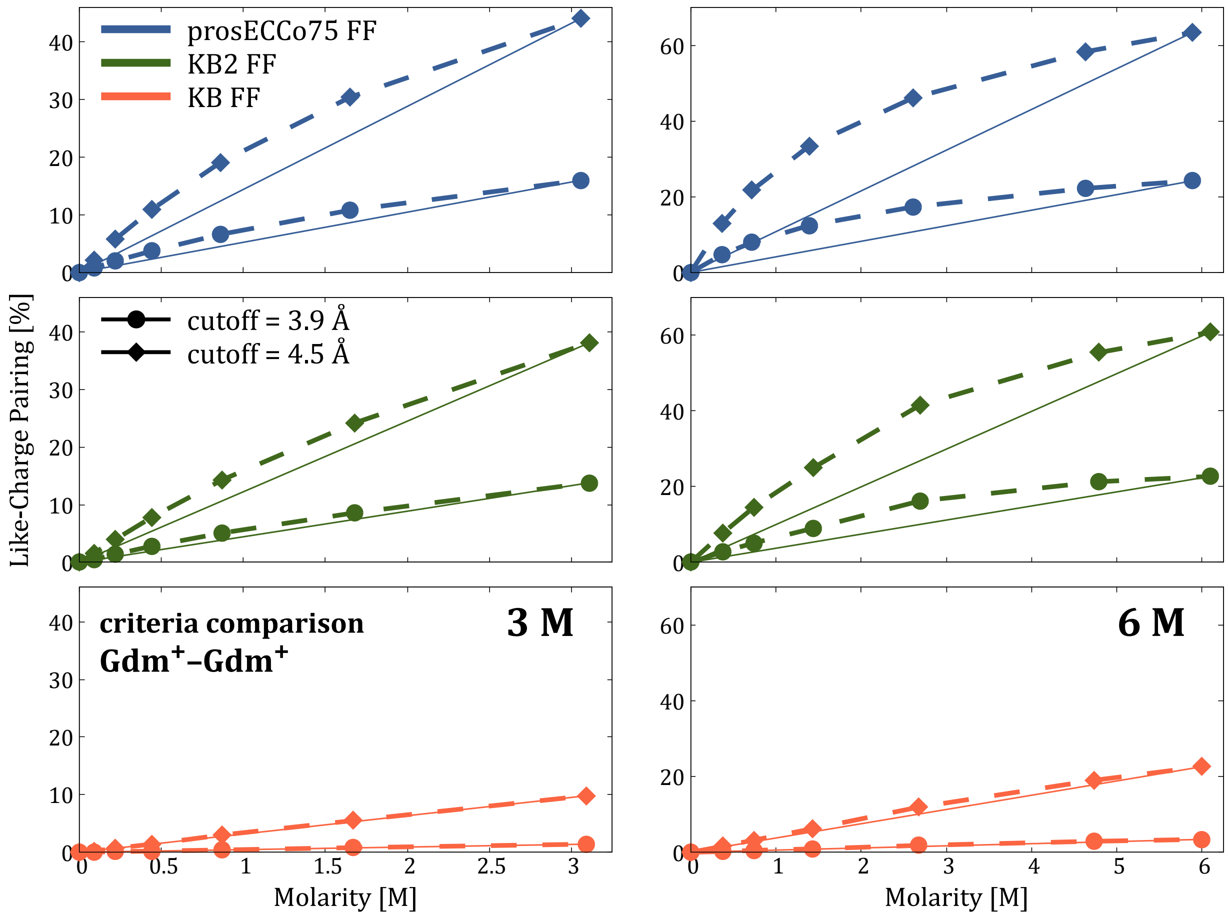


**Figure S8:** The ion pairing of the $Gdm^+$ cations as a function of $Gdm^+$ concentration in an aqueous solution, where a constant ionic concentration of either ∼3 M or ∼6 M is maintained by adding NaCl.

### CryoEM Experiments

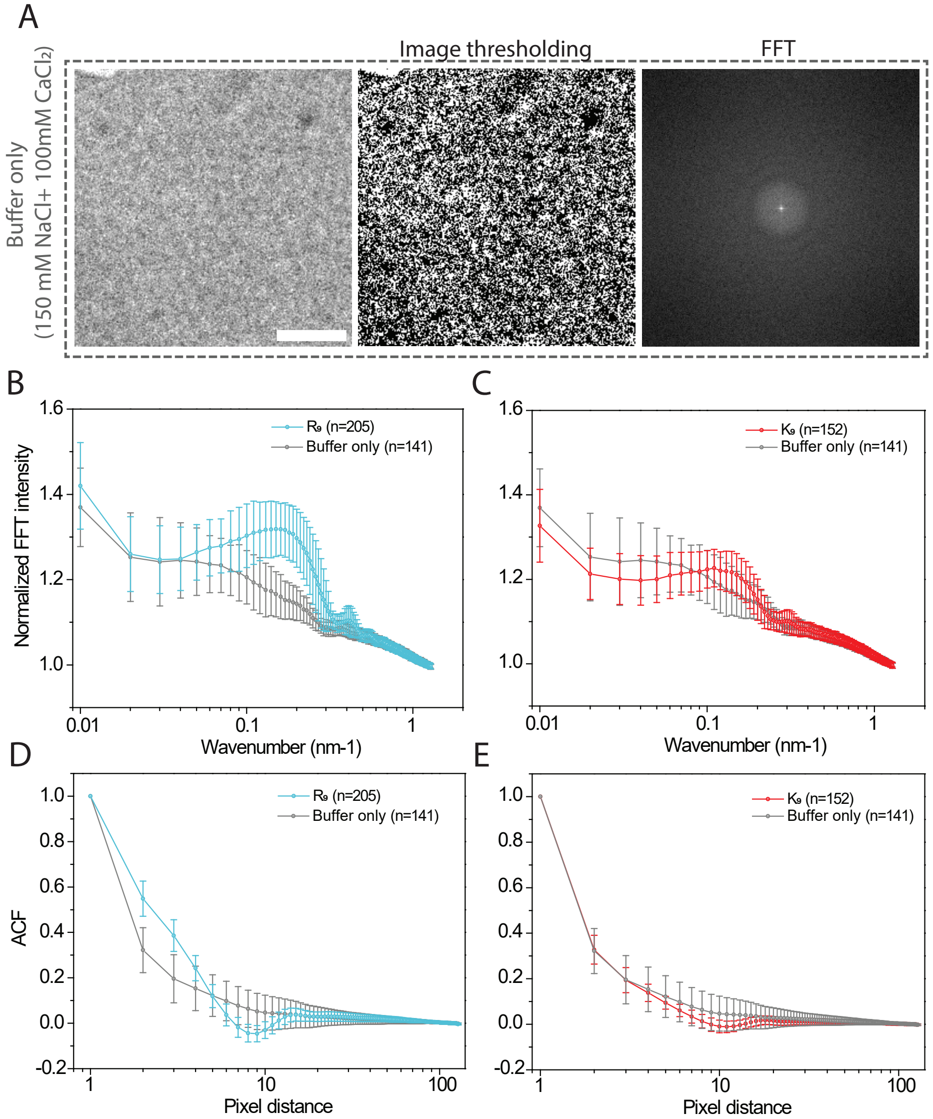


**Figure S9:** (A) Representative cryo-EM images of 150 mM NaCl and 100 mM $CaCl_2$ buffer, along with the corresponding binarized image used for autocorrelation function (ACF) calculation and Fast Fourier Transform (FFT) power spectra derivation. The scale bar for cryo-EM image is 50 nm. (B) Comparison of radial profiles derived from FFT power spectra images of 100 mM $R_9$ (blue) and buffer-only samples (gray). (C) Comparison of radial profiles derived from FFT power spectra images of 100 mM $K_9$ (red) and buffer-only samples (gray). (D) Comparison of spatial ACFs measured from cryo-EM images of 100 mM $R_9$ (blue) and buffer-only solutions (gray). (E) Comparison of spatial ACFs measured from cryo-EM images of 100 mM $K_9$ (red) and buffer-only solutions (gray). Data are presented as mean ± standard deviation, based on n = 205 frames for $R_9$, n = 152 frames for $K_9$, and n = 141 frames for buffer-only samples.
